# Growth characterization and biostimulant potential of *Coelastrella* sp. D14 a green microalga isolated from a solar panel

**DOI:** 10.1101/2024.01.24.576708

**Authors:** Sara Baldanta, Alice Ferreira, Luisa Gouveia, Juana Maria Navarro Llorens, Govinda Guevara

## Abstract

Extremophile environments are an important source for finding microorganisms with a potential for biotechnological applications. Among these microorganisms, microalgae contribute to several sustainable processes such as wastewater treatments or nutrition. In this work it was characterized a microalga isolated from a solar panel. The morphological and phylogenetic analysis revealed that the isolate collected was a *Coelastrella* strain. Cultivation and stress experiments has shown that *Coelastrella sp.* D14 can resist a long period of desiccation and it can grow on cheap sources such as piggery wastewaters (PWW). This work reports that a *Coelastrella* strain displays biostimulant properties with a germination index of 123% on *Lepidium sativum* when D14 biomass grown at 10% piggery effluent was used. Altogether, these results suggest that this novel strain could be a good chassis for further biotechnological applications.

**Highlights:**

- *Coelastrella* sp. D14, a xero-tolerant strain, has been isolated from a solar panel
- This strain can grow on piggery wastewater
- *Coelastrella* sp. D14 can promote germination of *Lepidium sativum*

## 1. Introduction

Water scarcity and pollution, recognized as significant environmental concerns, have garnered widespread attention, prompting efforts towards finding solutions (Hussain et al., 2021). Microalgae are one of the most attractive biological agents to address the water pollution problems and greenhouse effect. Microalgae are a highly diversified group of photosynthetic microorganisms adapted to a wide range of ecological habitats that utilize solar energy to generate biomass. Among the advantages of its use, it stands out their low nutritional needs, not depending on arable land nor potable water, they can grow under several stresses and can be harvesting daily (Rizwan et al., 2018; Tang et al., 2020). Additionally, various microalgal species have demonstrated the ability to thrive in municipal and/or industrial wastewaters, effectively eliminating organic carbon, nitrogen, and phosphorus, while fixing CO_2_. Moreover, many industries used microalgal feedstocks to get high-value co-products from the biomass such as antioxidants, lipids, vitamins, pigments, or carbohydrates besides biofuel to improve the economics (Sudhakar et al., 2019; Nayana et al., 2022).

Microalgal growth depends on both chemical and physical factors such as the type and concentration of carbon sources and minerals present in the medium, light intensity and regime (dark/light), pH, agitation, or temperature (Singh et al., 2015). For instance, a shortage in nitrogen or phosphorus alters the biochemical composition of the microalgae but also causes a drop in the growth rate (Procházková et al., 2014). Similarly, physical parameters affect the biomass production depending on the microbial species (Daneshvar et al., 2021; Elisabeth et al., 2021; Khanra et al., 2021). The ability of microalgae to acclimate to demanding wastewater conditions and endure oxidative stress particular to these environments differ among species. However, minimizing the cost of biomass production must be considered, and therefore, an equilibrium between growth and the use of cheap media must be reached. This strategy allows both i) wastewater remediation by recovering nutrients and removing pollutants from the environment and ii) the use of the biomass produced for different applications such as biofertilizers, bioplastics, cosmetics or ingredients in functional foods and feeds (Ferreira et al., 2017; Posadas et al., 2017; Ferreira et al., 2018; García et al., 2018; Ferreira et al., 2019; Viegas et al., 2021).

The utilization of biofertilizers and biostimulants as a natural product is particularly crucial to avoid the use of chemicals that may lead to environmental contamination, namely in soil, water and affect the quality of the food produced. Numerous efforts are being made to expand the application of these natural biostimulants (Navarro-López et al., 2020; González-Pérez et al., 2021; Sánchez-Quintero et al., 2023) under strict legislations and regulations that depends on the continent (Su et al., 2023). The use of microalgae as biostimulants has acquired importance for their role in the sustainability and circular bieconomy agenda (Ajeng et al., 2022; Sánchez-Quintero et al., 2023). This is because they are capable of sequestering CO_2_, they can survive in challenging environments such as waste effluents and and their easier cultivation compared to macroalgae (Sánchez-Quintero et al., 2023).

The genus *Coelastrella* (Chlorophyta phylum, Sphaeropleales order, Scenedesmaceae family) are green microalga with reported applications for bioremediation and value-added products such as UV-protective compounds among others (Zaytseva et al., 2021). *Coelastrella* is also a better renewable energy resource feedstock with a total of 18% of their biomass made up of lipids beneficial for biodiesel conversion (Nayana et al., 2022). This genus is mainly unicellular, ellipsoidal cells with a peculiar apical wart-like wall thickenings (John, 2002; Wang et al., 2019; Goecke et al., 2020; Maltsev et al., 2021). It can be often found in subaerial and terrestrial habitats, and it is universally distributed from the arctic boreal zone to tropical zones (Nayana et al., 2022). The strains isolated from extremophilic environments display unique properties for biotechnological applications as bioprospection of extremophiles have discovered strains with high resistance to various stresses such as withstanding extreme dehydration, salt stress, and high light exposure. Some examples are: *Coelastrella thermophila* var. globulina isolated from an algerian hot spring produces n-6 and n-3 polyunsaturated fatty acid of commercial interest (Boutarfa et al., 2022); *Coelastrella terrestris* collected from red mucilage in a glacier foreland in Iceland is proposed for biotechnological adonixanthin production (Doppler et al., 2022); a *Coelastrella* sp. isolated from an ammonia-rich environment could process piggery wastewater while using its biomass for other purposes such as biodiesel (Lee et al., 2021).

One of these extreme environments of interest are solar panels, an extreme habitat subjected to different stresses, such as high irradiation, temperature fluctuations, and desiccation (Dorado-Morales et al., 2016; Porcar et al., 2018; Tanner et al., 2018). The present work describes the identification and characterization of a *Coelastrella* sp. D14 strain from an extreme environment, a solar panel. The biotechnological potential of this novel strain was evaluated: D14 resists long periods of desiccation, it can grow on cheap sources such as piggery wastewaters (PWW), and for the first time this work reports that a *Coelastrella* xerotolerant strain can be used as a biostimulant.

## 2. Material and Methods

### 2.1. Strain isolation and culture conditions

The microalga used in this study, *Coelastrella* sp. D14, was isolated from a solar panel in Valencia (Spain) (Baldanta et al., 2023). *Coelastrella* sp. D14 was grown in BG11 medium on 1.5% agar plates or liquid medium at 30°C ± 2 °C under 100 μE·m^−2^·s^−1^ of continuous white light, under orbital shaking (150 rpm). The BG11 medium contained 1.5 g/L NaNO_3_; 0.02 g/L Na_2_CO_3_; 0.03 g/L K_2_HPO_4_; 0.075 MgSO_4_ * 7 H_2_O; 0.036 g/L CaCl_2_ * 2 H_2_O; 1 g/L Na_2_EDTA * 2 H_2_O; 1.81 g/L MnCl_2_* 4 H_2_O; 0.05 g/L CoCl_2_* 6 H_2_O; 0.039 g/L Na_2_MoO_4_*H_2_O; 0.08 g/L CuSO_4_* 5 H_2_O; 0.22 g/L ZnSO_4_ * 7 H_2_O; 2.86 g/L H_3_BO_3_; 6 g/L citric acid and 6 g/L ferric ammonium citrate (PhytotechLabs) (Rippka et al., 1979). BG11 medium was buffered to pH 7.5 with 10 mM HEPES. The cultured algal cells were observed and photographed under a microscope (Leica, model DM750). Axenic strains were stored at −80°C in BG11 medium supplemented with 5% (v/v) DMSO.

Cultures of *Coelastrella* sp. D14 were harvested at different growth phases and preparations of these cultures were photographed and then processed by the LAS V4.2 software. ImageJ software was used to measure the cell size.

### 2.2. Molecular identification and Phylogenetic analysis

*Coelastrella* sp. D14 was previously identified using primers for 18rRNA amplification (18S-Fw 5’-GTCAGAGGTGAAATTCTTGGATTTA-3’, 18S-Rv 5’-AGGGCAGGGACGTAATCAACG-3’) (Baldanta et al., 2023). The 18S rRNA gene sequence of *Coelastrella* sp. D14 (PP158241) was searched against homology sequences in Genbank using BLAST (http://blast.ncbi.nlm.nih.gov). 18S rRNA gene sequence from the identified *Coelastrella* sp. D14 and the top BLAST sequences were used for phylogenetic analysis. Multiple sequence alignments were performed using MUSCLE algorithm in MEGA-X. A neighbor-joining phylogenetic tree was built with the aligned sequences based on the K2+G+I model with a bootstrap analysis involving 1000 resampling trees using MEGA-X package. *Dunaliella salina* 18S rRNA was used as outgroup.

### 2.3. Autotrophic growth

Axenic *Coelastrella* sp. D14 was inoculated into 20 mL of BG11 medium in 100 mL Erlenmeyer flasks to an initial optical density at 750 nm (OD_750nm_) of 0.05 and grown at different salinity conditions, pH, or nitrogen sources. To assess growth at different salt concentrations, BG11 medium was prepared containing 0.1, 0.25, 0.5, or 1 M of NaCl. The influence of pH on cyanobacterial growth was explored in BG11 buffered to pH 4, 6.5, 9, and 11 with 10 mM Tris adjusted to each pH. To examine the strains for growth on different nitrogen sources, BG11 was modified by replacing the 16 mM of NaNO_3_ with 16 mM of NH_4_Cl or urea. Tolerance to urea was determined by adding this compound to final concentrations of 8 and 16 mM to BG11. For temperature experiment tests, 100 mL Erlenmeyer flasks with an initial OD_750nm_ of 0.20 were used. As a control, strains grown under routine conditions (BG11 pH 7.5, 30°C, 150 rpm, and continuous light 100 μE·m^−2^·s^−1^) were used. The temperature effect on growth was evaluated at 4, 40, and 50°C, using 30°C as control keeping the other conditions constant. Cell growth was monitored by measuring the OD_750nm_ for a 10-day period. In all the growth experiments, three biological replicates were performed. To define the relationship between cell density per unit OD at 750 nm wavelength, a hemocytometer was used to count the cells. Growth rate was determined by plotting the log OD versus time and calculating the slope in the linear portion, related to the exponential growth. The beginning of the growth phase was considered when the growth of the cyanobacteria was appreciable. Doubling time corresponds to the log2/r. In addition, the biomass dry weight and the ash free dry weight (AFDW) were determined through gravimetry by drying the samples at 105°C overnight and incinerating at 550°C for 1 h, respectively.

### 2.4. Heterotrophic and Mixotrophic Growth

First, to assess the heterotrophic growth of *Coelastrella* sp. D14, BG11 agar plates were prepared at final concentrations of 10 mM with different carbon sources: glucose, sucrose, lactose, arabinose, maltose, fructose, galactose, mannose, and glycerol. The tests were performed with spots of 10 μL at OD_750nm_=1 onto BG11 plates to reduce the possibility of contamination. Plates were incubated at 30°C in darkness for 30 days. Furthermore, the photosynthesis inhibitor DCMU (3-(3,4-dichlorophenyl)-1,1-dimethylurea) was added for a final concentration of 10 μM to make sure that the observed growth was heterotrophic. The cell growth was evaluated by checking the appearance of colonies after the incubation period.

Once the sugars were determined for *Coelastrella* sp. D14 cultivation, the heterotrophic and mixotrophic growth were evaluated in liquid medium. Axenic *Coelastrella* sp. D14 was inoculated into 20 mL of BG11 medium in 100 mL Erlenmeyer flasks to an initial optical density at 750 nm (OD_750nm_) of 0.3 and grown in light conditions with no sugar, and glucose or mannose at 10 mM (mixotrophic growth). In parallel, the same conditions were used adding the photosynthesis inhibitor DCMU at 10 μM (heterotrophic growth). Cell growth was monitored by measuring the OD_750nm_ for a 7-day period. In all the growth experiments, three biological replicates were performed. Cultures were checked to ensure that they were free of contaminant bacteria before the experiments.

### 2.5. Desiccation-Tolerance Test

Microalgae strain was grown on 9–10 mL of BG11 agar plates (6 cm diameter) under continuous light (60-80 μE·m^−2^·s^−1^) at 30°C for 2 weeks at 30–35% relative humidity. Then, plates were left to be air-dried under routine growth conditions by removing the parafilm from the Petri dishes. After about 15 days, dried cultures were stored in the laboratory bench at room temperature for 3 months, 7 months, and 1 year. For the 1-year dried samples, some samples were maintained in parallel under routine growth conditions. For rehydration, the dried samples were soaked with 1 mL of sterile water for 15 min at room light, streaked on BG11 plates, and incubated under the same initial conditions (60–80 μE·m^−2^·s^−1^ and 30°C). Results were observed after 2–3 weeks. As a negative control for desiccation tolerance, *Synechocystis* sp. PCC 6803 was used.

### 2.6. Wastewater treatment and biomass production

#### 2.6.1. Wastewater characterization

The piggery wastewater (PWW) was collected from a stabilization pond in a local pig farm from Valorgado in Herdade do Pessegueiro (39◦00009.000 N, 8◦38045.500 W) (Glória do Ribatejo, Portugal). This effluent corresponds to the liquid fraction of pig slurry after separation (sieve 1-10mm) from solid manure. The nutrient composition of PWW was determined by standard methods. The Kjeldahl nitrogen (TKN) was determined by a modified Kjeldahl method adapted from the standard method 4500-Norg B (Clesceri et al., 1988). Ammonium nitrogen was quantified by titration after a distillation step based on standard methods 4500-NH 3 B and C (Clesceri et al., 1988). A commercial kit was used for the measurement of phosphorus (Phosver 3-Powder Pillows, Cat. 2125-99, HACH) using a HACH DR/2010 spectrophotometer, at 890 nm. COD determination was carried out according to the open reflux method—Method 5220-B. The effluent composition is shown in Table 1.

**Table 1.** Composition of piggery wastewater: pH, total Kjeldahl nitrogen (TKN), ammonia (NH_4_^+^), phosphate (PO_4_^3-^), and chemical oxygen demand (COD).

| pH | TKN (mg N/L) | $\text{NH}_4^+$ (mg/L) | $\text{PO}_4^{3-}$ (mg/L) | COD (mg $\text{O}_2$/L) |
| --- | --- | --- | --- | --- |
| 7.72 | 1333±6 | 1281±1.4 | 218±5 | 4396±94 |

#### 2.6.2. Microalga screening

A *screening* was carried out to determine if D14 was able to grow in PWW. 50 mL flasks were inoculated using 20 mL with different dilutions (1:20, 1:10, 1:5, and 1:2.5) of PWW with tap water as the cultivation medium and were kept at room temperature, under at light intensity of 41 μE·m^−2^·s^−1^, and orbital agitation at 150 rpm in an incubator shaker (New Brunswick Scientific Co, USA).

#### 2.6.3. Biomass production

To obtain biomass, the microalga cultures were cultivated in 1 L bubble columns photobioreactors (PBRs) using 1:20 (PWW) or BG11 as medium. The cultures were maintained at room temperature (23-25°C) under continuous illumination at an average light intensity of 60 μE·m^−2^·s^−1^. The aeration was supplied at 0.15 vvm (air volume (L) per volume of culture medium (L) per minute (m) from aquarium pumps.

The microalga cultures were cultivated in 5 L bubble columns photobioreactors (PBRs) using the same 1:20 and 1:10 PWW as medium, at a working volume of 1 L. The cultures were at room temperature (23-25°C) under continuous illumination (3 fluorescent lamps of 36 W and 6 of 18 W, Philips TL-D) at an average light intensity of 53 μE·m^−2^·s^−1^. The aeration was supplied at 0.15 vvm (air volume (L) per volume of culture medium (L) per minute (m) using aquarium pumps. After 9 days of cultivation the biomass was collected by centrifugation (10.000×g, 10 min).

### 2.7. Biomass characterization

The biochemical composition of the microalgal biomass was determined in terms of proteins, carbohydrates, lipids, moisture, and ash. All analyses were performed in triplicate. The protein content was determined following the method described by (González López et al., 2010), which is a modification of the Lowry method, with BSA (Bovine serum albumin) as the standard. The sugar content was determined by the phenol-sulfuric method (Dubois et al., 1956) after quantitative acid hydrolysis (Hoebler et al., 1989) of the biomass. A calibration curve was prepared using standard glucose solutions. Lipid content was determined gravimetrically after Soxhlet extraction with n-hexane during 6h, using biomass previously submitted to bead milling (Retsch MM400, 25 Hz for 3 min and 50 seconds).

### 2.8. Germination Index

The biostimulant activity of the microalga *Coestrella* sp. D14 was determined by measuring the germination index of seeds of *Lepidium sativum*, according to the method described by (Zucconi et al., 1981).

Microalga culture (whole biomass) and extracts obtained from the growth at different conditions (BG11, 5%, 10%, and 20% PWW) were tested at different concentrations (0.1, 0.5, 1, and 2 g/L). Microalga extracts were prepared by submitting the harvested biomass to high-pressure homogenization (1200 bar for 1 cycle) to disrupt the cells (Ferreira et al., 2022). Treatment solutions with microalga culture and extracts were then prepared with distilled water to the desired concentrations. A total of 32 treatments were tested (Fig. 1).

**Fig. 1.**
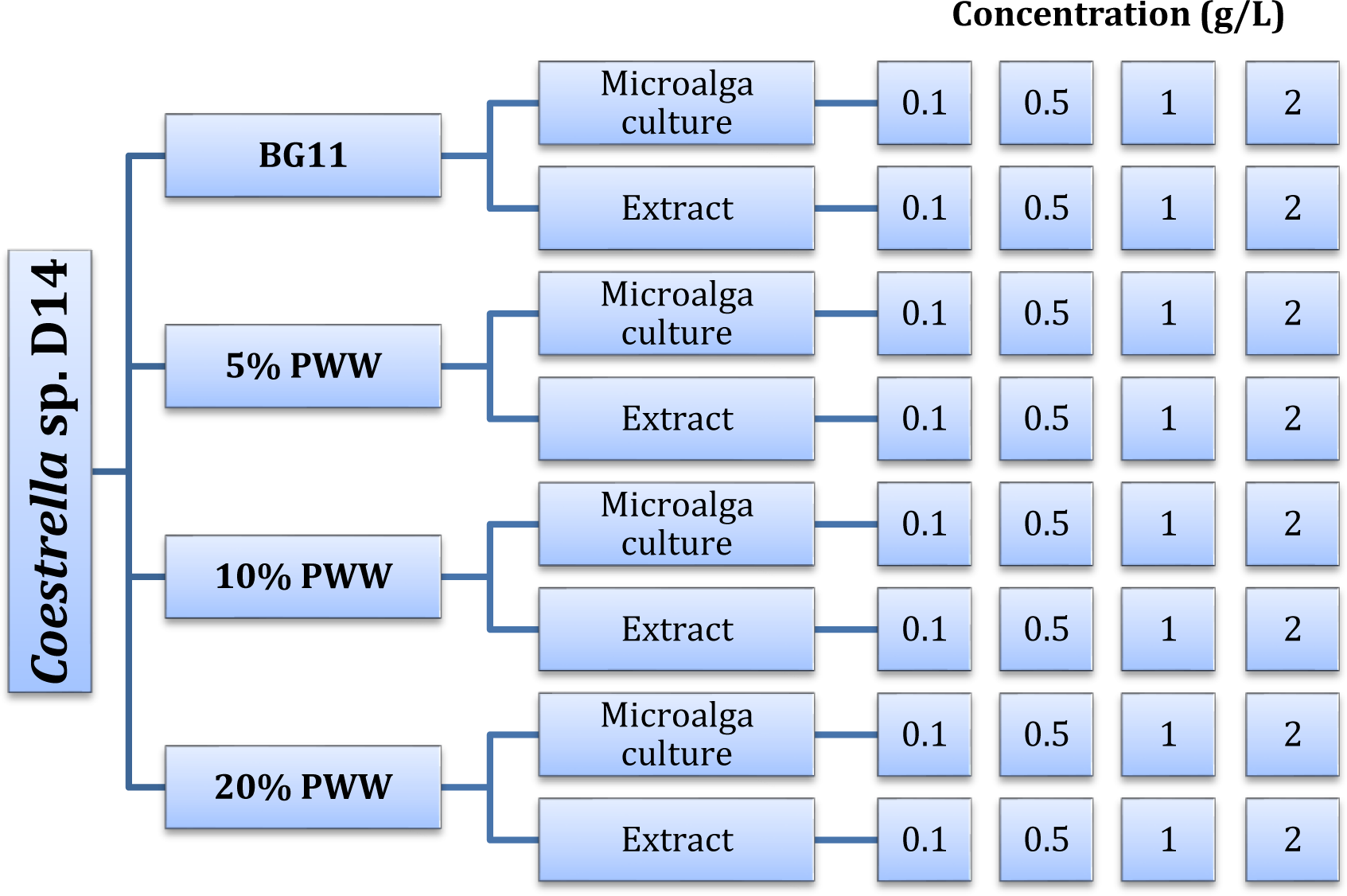
Schematic diagram of the treatments of *Coestrella* sp. D14 tested in the germination trials. Different growth media (BG11, 5, 10, and 20% PWW), different biomass processing (microalga culture and extract from disrupted biomass), and different treatment concentrations (0.1, 0.5, 1, and 2 g/L) were tested.

The germination experiments were carried out in sterilized rectangular Petri dishes (10 mm x 17 mm) with Whatman No 5 filter papers wetted with 7 mL of each treatment solution, with 10 seeds per dish in duplicates. Distilled water was used as the negative control. All samples were incubated at room temperature (25 °C) in the dark for 3 days and the Petri dishes in a vertical position. At the end of 3 days, the seedlings were photographed and measured with the program ImageJ (Rasband, 1997). Results were registered for comparison between the microalga treatments and the control with distilled water.

Finally, the germination index was determined by the Equation (1), where G and L are the number of germinated seeds and their length in the case of the microalgal cultures and Gw and Lw are the same parameters but in the control (distilled water). The data shown in the germination index experiments is, therefore, the result of the measurement of 100 seeds for each treatment.

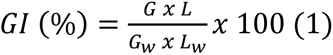

### 2.9. Statistical Analyses

One-way Anova with post-hoc Tukey HSD, with Scheffé, Bonferroni and Holm multiple comparison results were calculated in the different conditions studied in this work on Astatsa.com (https://astatsa.com/; [Vasavada, 2016](https://www.frontiersin.org/articles/10.3389/fmicb.2023.982422/full#ref86)). Correlation was considered statistically significant when p < 0.05.

## 3. Results

### 3.1. Isolation and identification of *Coelastrella* sp. D14

Solar panel samples were collected during the summertime of 2013 and 2014 for screening cyanobacteria and microalgae. The first isolation was made in Castenholz-D medium (Baldanta et al., 2023) and afterwards, microalgae and cyanobacteria were maintained growing on BG11. At this time, a consortium among microorganisms, bacteria and cyanobacteria/microalgae, was evident on BG11 plates. After several streaks, different strains were isolated (Baldanta et al., 2023). One of these strains, a unicellular green microalga, named D14, was identified by PCR 18S rRNA amplification reaching a homology of 99% with other *Coelastrella* strains.

A phylogenetic analysis based on the 18S rRNA gene sequence and a comparison to similar strains in the GenBank database indicated that the D14 had a high similarity with other strain sequences of *Coelastrella* (Fig. 2).

**Fig. 2.**
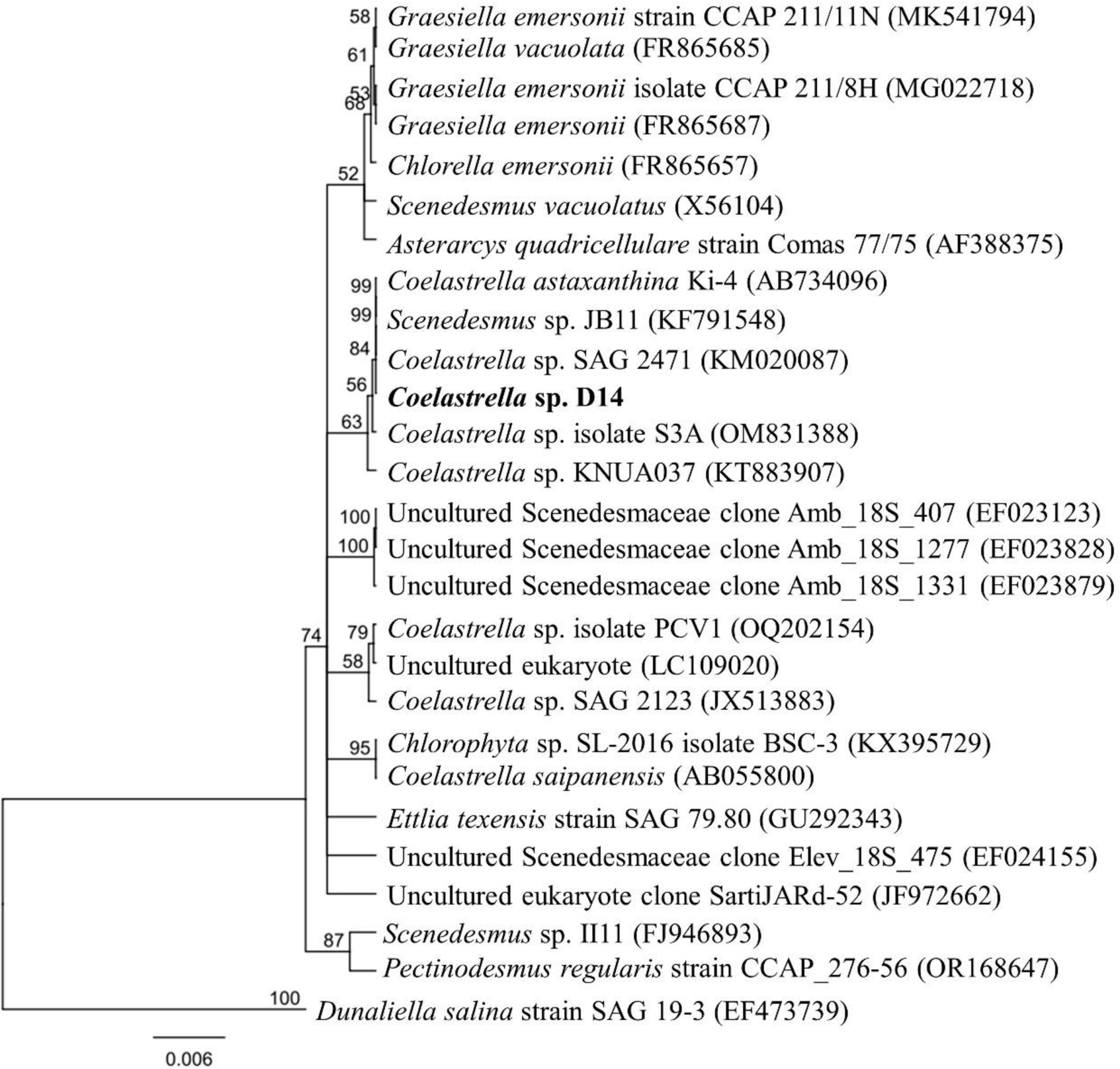
Phylogenetic tree inferred from 18S rRNA gene sequence from *Coelastrella* sp. D14 isolated from the solar panel. The tree was constructed using the neighbor-joining method in MEGA-X. The length of the scale bar indicates 0.006 substitutions per site. The percentages of bootstrap support of branches (>50%) are indicated at each node. The numbers in the parenthesis are the accession numbers of respective 18S rRNA gene sequences obtained from Genbank. *Coelastrella* sp. D14 is shown in bold.

Streaked microalgal colonies on agar plate, and microscopic observation is shown in Fig. 3 (A-H). Light microscopical observations showed that *Coelastrella* sp. D14 was unicellular green coccoid microalgae. The cells showed usually as single oval cells, but a large degree of variation in cell sizes was observed ranging between 4.2 to 14.8 µm, with a mean diameter of 8.68 ± 1.96 μm. Single cells are smaller, have a lemon-shaped after division, have a thin wall, and the pyrenoid is clearly noted. Also, in some of them, wart-like wall thickenings are observed (Fig. 3E). As the culture grew, the cells appeared round shaped and formed small groups of 2-6 cells. Fig. 3G shows 2-3 daughter cells after cell division with the cell wall of the mother cell surrounding the new cells. In mature cells, the chloroplast is dissected into blades.

**Fig. 3.**
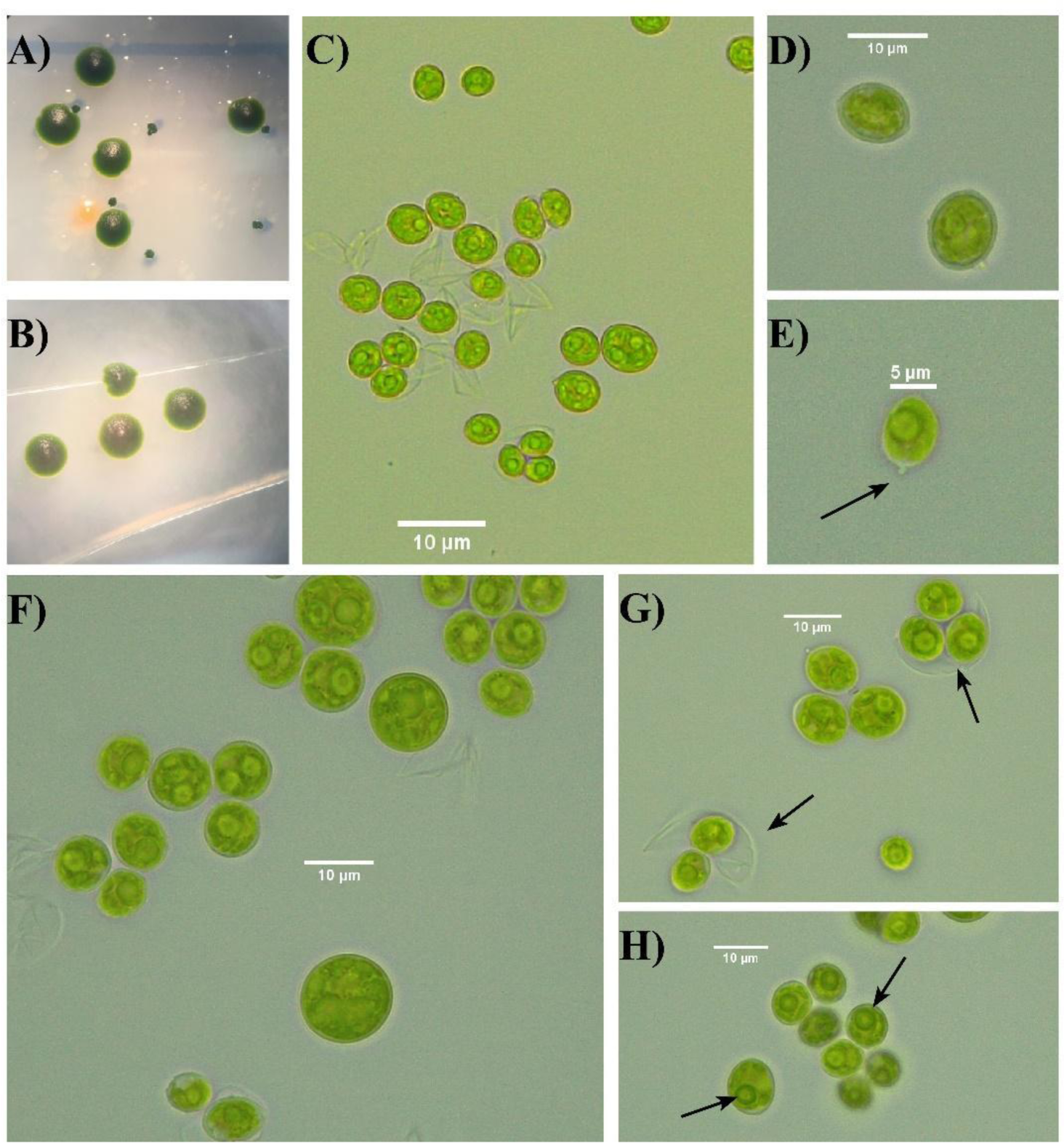
Isolation of *Coelastrella* sp. D14. **A)** Streak colonies from the original consortium microalgae-bacteria. **B)** Streak colonies from the axenic microalgae. **D-H)** Bright-field photomicrographs of *Coelastrella* sp. D14 on different days of culture. Scale bar is shown in all the pictures. D and E, lemon-shaped cells, arrow indicates a wart-like wall thickening. F, mature and round-shaped cells. G and H, cells after division. In G arrows indicate the cell wall of the mother cell and in H, the pyrenoids are pointed.

### 3.2. Autotrophic growth conditions for *Coelastrella s*p. D14

The growth evolution of *Coelastrella* sp. D14 in BG11 medium (pH 7.5) within 10 days at 30°C under 100 μE·m^−2^·s^−1^ were studied (Fig. 4, Table S1). A mild lag phase of two days was observed and then, the microalgae reached 2.60 ± 0.01 days of doubling time. The calculated correlation between the OD_750nm_ and cells per mL was: N°cells/mL = 3.191*10^6^ OD_750_ (R^2^ = 0.99). In addition, the correlation of the biomass dry weight with OD_750nm_ was determined through gravimetry (Dry weight=0.5314* OD_750_-0.0127; R^2^ = 0.99). After 10 days of growth in BG11, a total of 2.53 g/L of *Coelastrella* sp. D14 biomass was obtained.

**Fig. 4.**
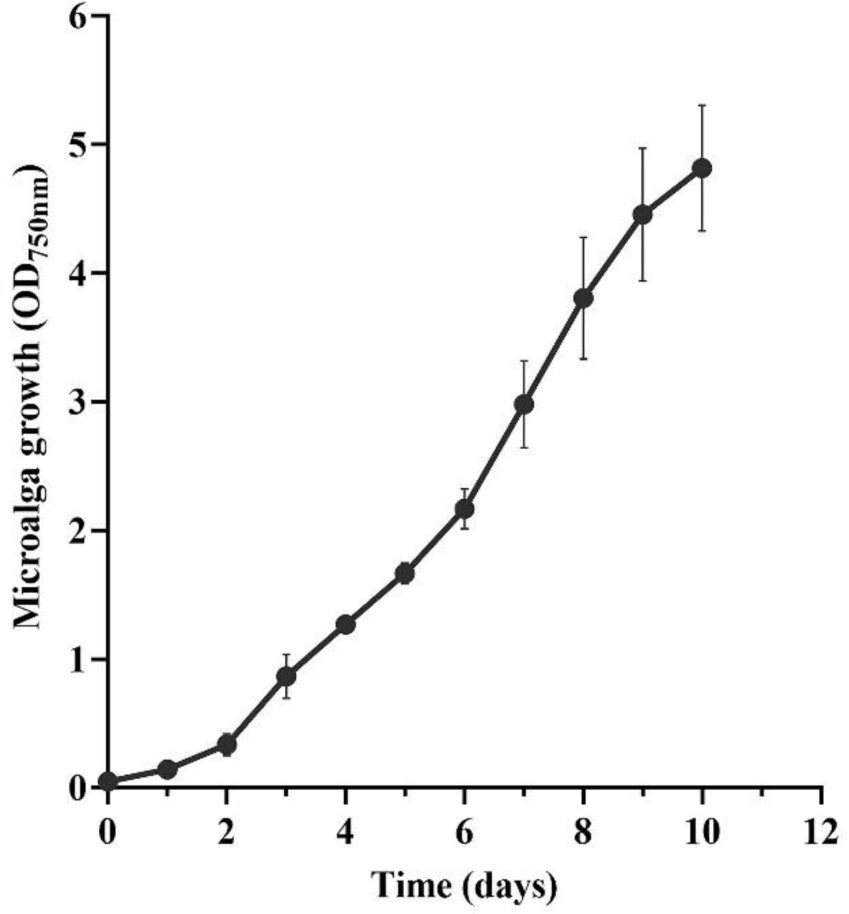
Growth curve of *Coelastrella* sp. D14 in BG11. Average OD_750nm_ of three biological replicates together with the standard deviation.

Regarding salt stress, *Coelastrella* sp. D14 grew up to 0.5 M of NaCl (Fig. 5A). The growth was slightly affected at 0.1 M and 0.25 M of NaCl, as the statistical analysis revealed (Table S1). At 0.5 M of NaCl, the lag phase was 3 days longer and the maximal OD was lower compared to control. However, the doubling time was comparable to the control. No growth was observed at 1 M NaCl.

**Fig. 5.**
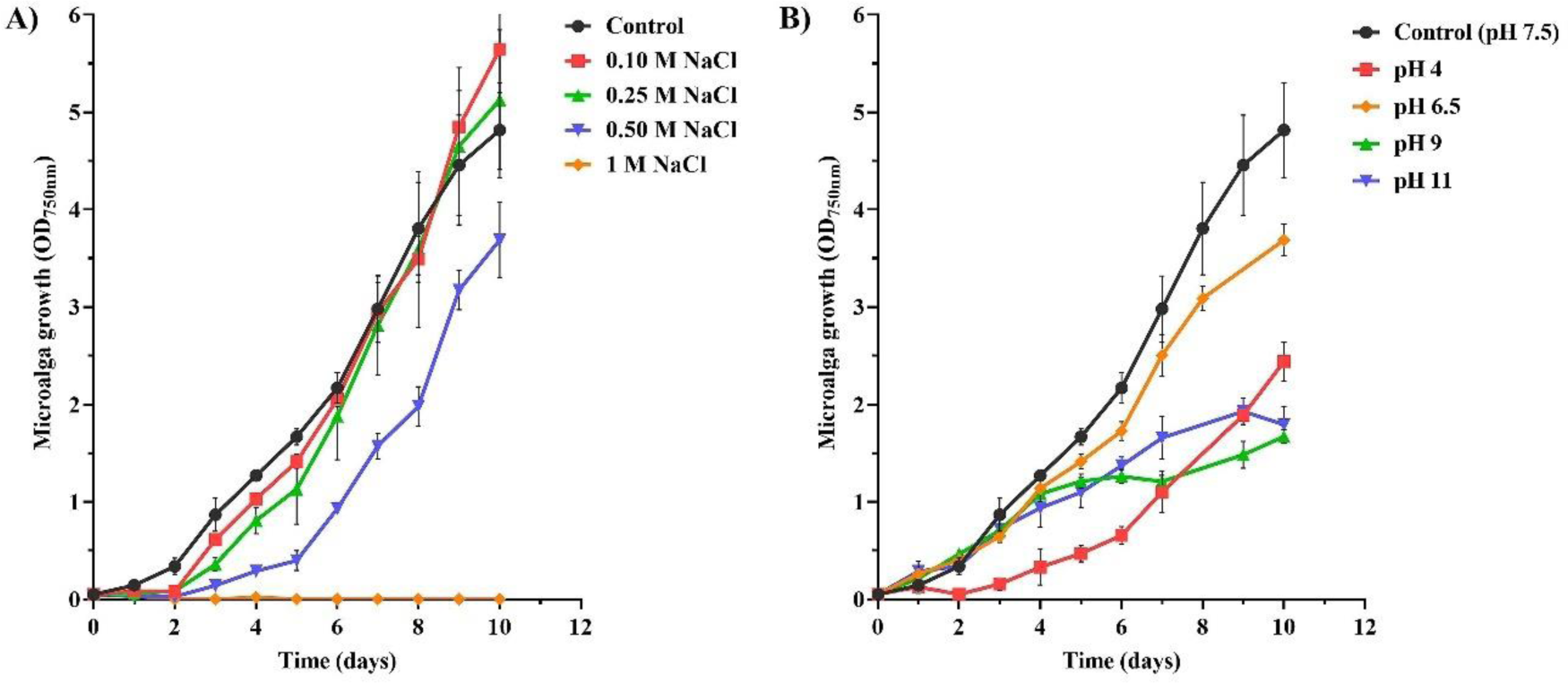
Growth characterization of *Coelastrella* sp. D14 in different conditions. **A)** Effect of NaCl concentration on microalgae growth. As a control, growth in BG11 medium was used. **B)** Effect of pH on microalga growth. BG11 buffered with 10mM Tris pH 4, pH 6.5, pH 9, and pH 11 was used instead of 10 mM HEPES at pH 7.5 (control). In all cases, the graphs show the average OD_750nm_ of three biological replicates together with the standard deviation (n=3).

The best pH to grow *Coelastrella* sp. D14 was 7.5 (control) as it is shown in Fig. 5B. non-statistical differences were observed in the doubling time for pH 6.5 when compared to the control; however, a lower growth was reached after 10 days. Interestingly, *Coelastrella* sp. D14 grown at pH 4 despite having an extended lag phase, the doubling time was similar to that calculated for pH 7.5, after 5 days of grown. On the other hand, the microalga was able to tolerate a pH 9 and 11, with a statistically significant higher doubling time and reaching a lower OD than the control (5.91 and 3.39 days, respectively vs. 2.57 days at pH 7.5, Table S1). The results demonstrate that *Coelastrella* sp. D14 is pH broadly resistant (from 4 to 11). More specifically, in those cultures, the final pH was ∼6.3 and ∼9.5, respectively and the cultures remained green after the experiment (Fig. S1).

Temperature greatly affected the growth of *Coelastrella* sp. D14 (Fig. S2). No growth was observed at 50°C, and the cultures were bleached on the second day of the experiment. At 4 and 40°C, we had a small increase in the microalga growth, and the cultures remained pale green, indicating they were highly stressed.

The effect of the absence or the use of other nitrogen sources at 16 mM (the same concentration as NaNO_3_ is used in BG11) for the cell growth was assessed in liquid cultures. Fig. 6A shows the growth of the microalga for 10 days using alternative nitrogen sources, such as urea or ammonium. *Coelastrella* sp. D14 was not able to grow without any nitrogen source, which may be because they lacked the ability to fix atmospheric nitrogen in the conditions tested. Growth in the presence of ammonium was also closer to the result observed in BG11_0_, despite an initial increase of the OD after 6 days. However, even though the microalgae displayed the best growth with nitrate, it seems that urea could also be used as a nitrogen source, with no statistical differences and with a similar doubling time (2.52 days) compared to the control but with a longer lag phase (4 days instead of 2 days).

**Fig. 6.**
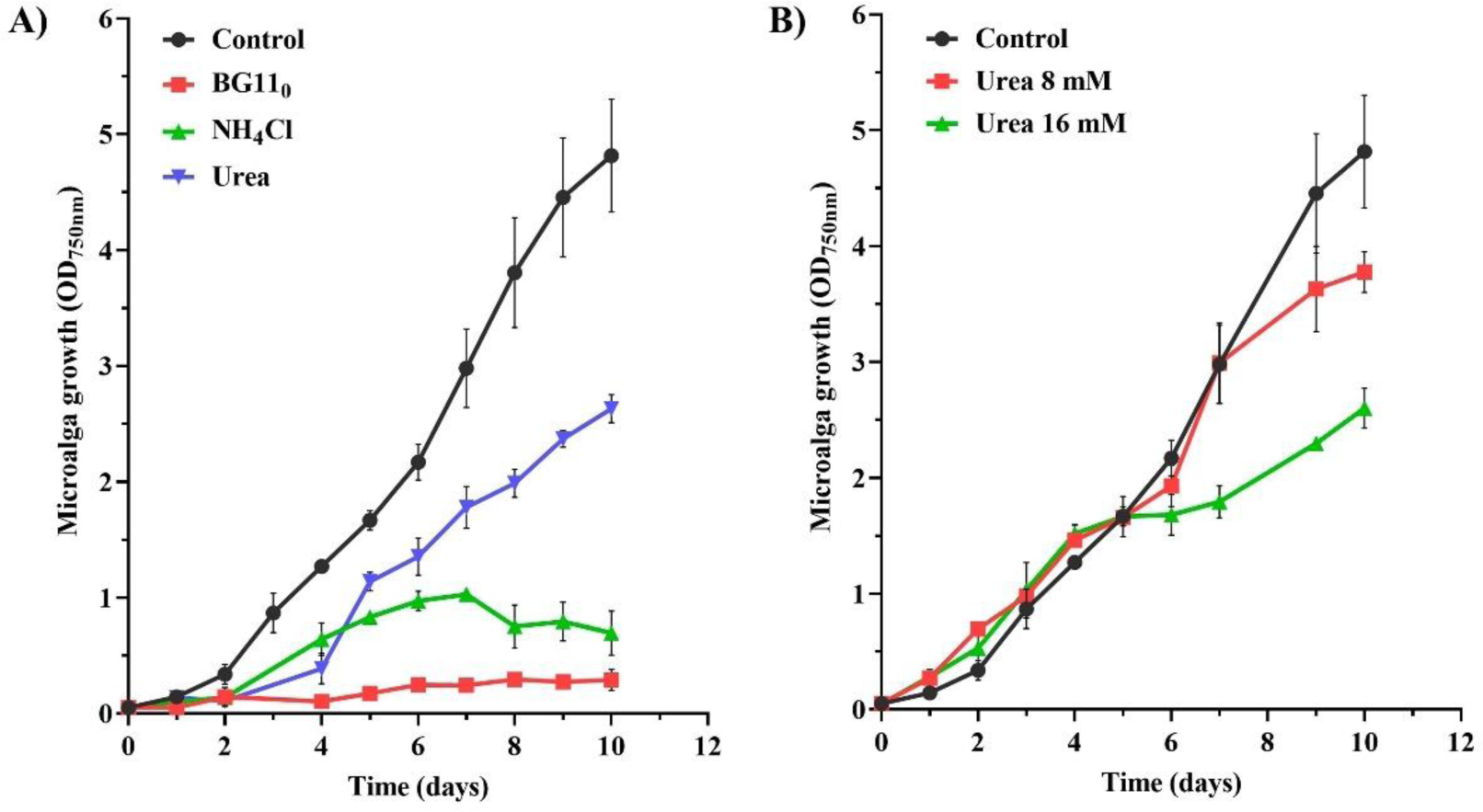
Effect of nitrogen source on 10 days growth of *Coelastrella* sp. D14. **A)** NaNO_3_ in BG11 was substituted for urea or NH_4_Cl (16 mM). Alternatively, no nitrogen was added (BG11_0_). **B)** *Coelastrella* sp. D14 tolerance to urea. Urea at 8 and 16 mM was added to BG11 complete medium. As a control, strains grown under routine conditions (BG11 pH 7.5, 30°C, 150 rpm, and continuous light 100 μE·m^−2^·s^−1^) were used. The graphs show the average OD_750nm_ of three biological replicates together with the standard deviation (n=3).

Considering that urea was the best nitrogen source for growing the microalga (Fig. 6A) and can be abundant in different types of wastewaters, various concentrations of urea were tested to grow *Coelastrella* sp. D14 (Fig. 6B). The microalga was sensitive to 16 mM urea, while 8 mM allowed a better growth, although lower than the control after 10 days of growth. These results show that even though urea cannot replace NaNO_3_ as a nitrogen source, some strains can tolerate it.

### 3.3. Heterotrophic and Mixotrophic Growth

The growth of *Coelastrella* sp. D14 was tested heterotrophically, as this strategy is widely used to increase microalgae biomass. First, it was valuated growth using glycerol and eight different sugars at 10 mM in BG11 agarized medium, complete darkness and with the photosynthesis inhibitor DCMU at 10 μM final concentration, for 30 days. The microalga was able to use glucose, fructose, and mannose as carbon source. *Coelastrella* sp. D14 showed a weak growth using sucrose but could not grow on maltose or lactose (Fig. 7A). Erlenmeyer flask experiments were done to find accelerated mixotrophic biomass growth compared to an autotrophic condition. The investigated C-sources were glucose and mannose as they contain the same amount of carbon (both are hexoses). In parallel, as control, heterotrophic growth was performed by adding DCMU to cultures with carbon source and light, to ensure the growth was due to the consumption of the sugar. Cultures without DCMU and carbon source represent the phototrophic growth.

**Fig. 7.**
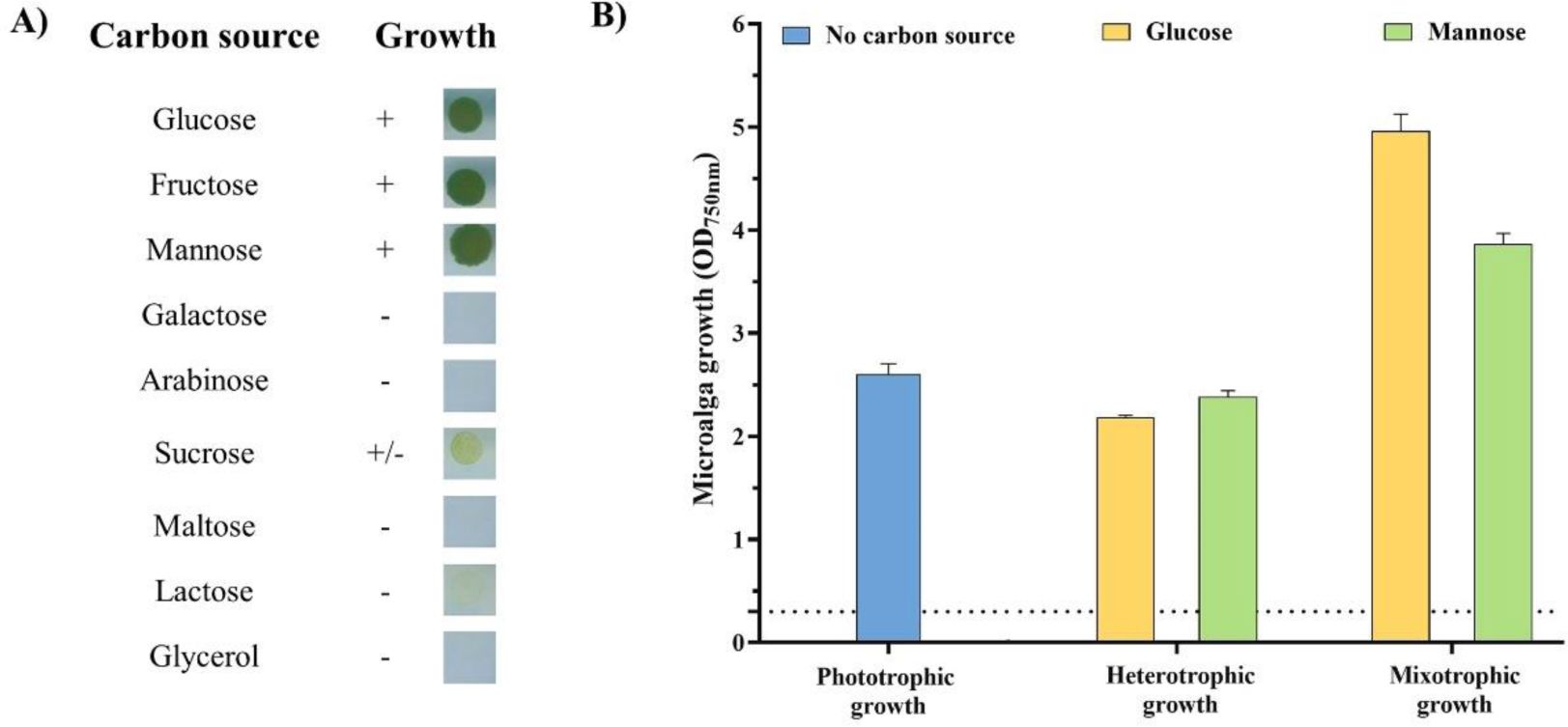
Growth of *Coelastrella* sp. D14 using different carbon sources. **A)** Heterotrophic growth of *Coelastrella* sp. D14 in agar BG11. Microalga was grown to OD_750nm_ = 1, 10 µL were plated and let the microalga grow for a month. **B)** Evaluation of phototrophic, heterotrophic and mixotrophic growth in *Coelastrella* sp. D14. For heterotrophic growth, the photosynthesis inhibitor DCMU was used at 10 µM concentration. Sugars were added at 10 mM. The circle crossed indicates that no sugar was added. OD_750nm_ was measured after 5 days of growth from an initial OD_750nm_ of 0.3 (dashed line). In all cases, the graphs show the average OD_750nm_ of three biological replicates together with the standard deviation (n=3).

As it is shown in Fig. 7B, the culture supplemented with glucose reached the highest OD_750nm_ compared to mannose (mixotrophic) and phototrophic growth. No differences were observed between the autotrophic control and the heterotrophic growth (DCMU condition), for both carbon sources.

### 3.4. Resistance of *Coelastrella* sp. D14 to desiccation

Considering that the microalga has been isolated from a solar panel, it was of interest to study its capacity to resist desiccation, so it was tested the desiccation-tolerance of *Coelastrella* sp. D14 for 3 months, 7 months, and 1 year. As shown in Fig. 8A, in terms of ability to form colonies and grow, the rewetted samples differed little from the non-dried cultivated form, indicating that *Coelastrella* sp. D14 was drought-resistant. *Synechocystis* sp. PCC 6803, which was used as a non-resistant strain, was unable to grow after 3 months, the minimum time tested (data not shown). Just after the rewetting, all the cells exhibited the same morphology, with a thick sheath surrounding the cells (Fig. 8D).

**Fig. 8.**
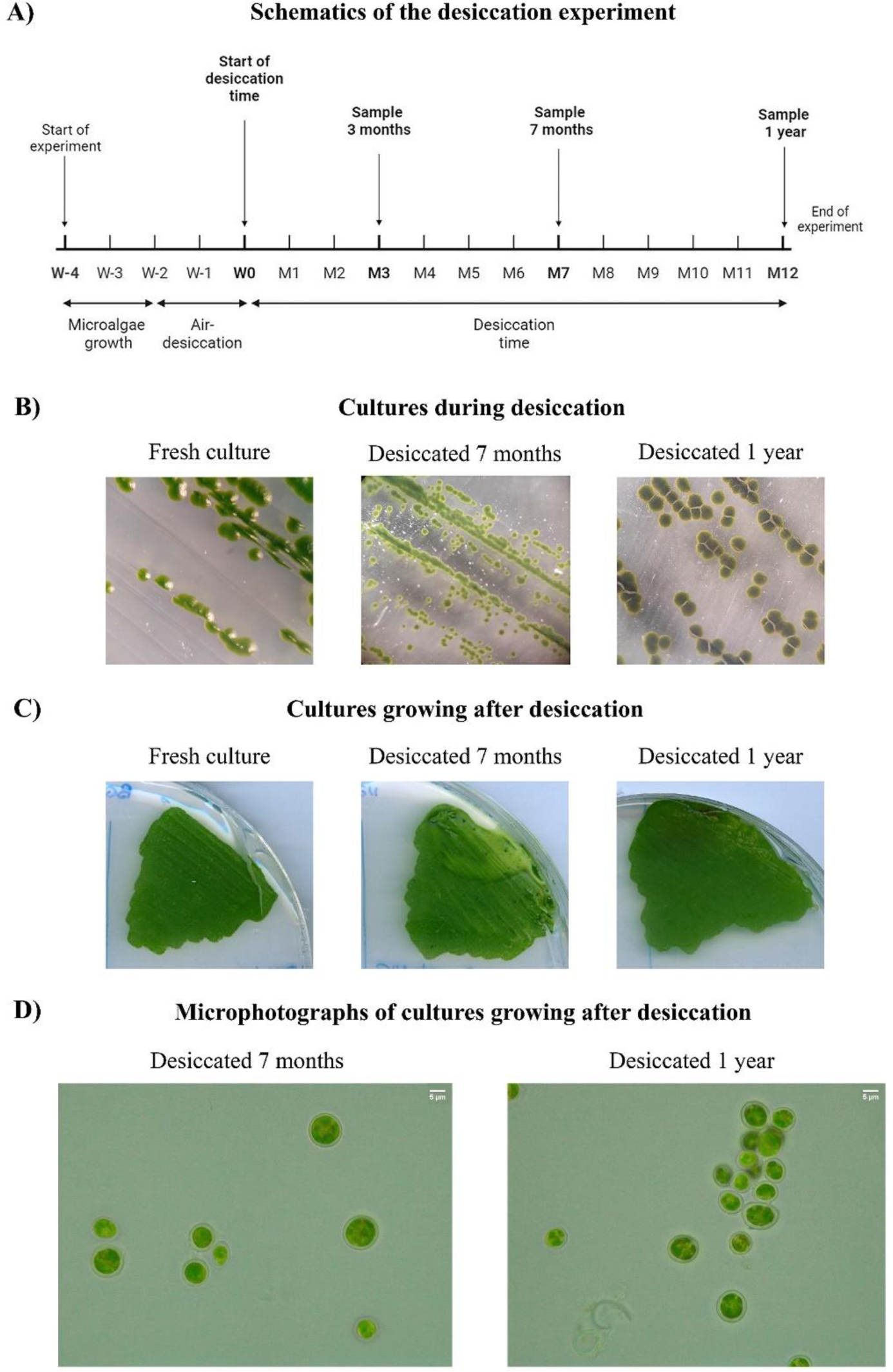
Long-term desiccation tolerance in *Coelastrella* sp. D14. **A)** Diagram of the desiccation process. W represents weeks, M represents months. **B)** Microphotographs of *Coelastrella* sp. D14 during the experiment. **C)** Growth observed after 2 weeks on BG11 post rewetting for 7-month and 1-year desiccated culture; a freshly streaked strain was included as a control. **D)** Light microscopical observation of *Coelastrella* sp. D14 just after rewetting 7 months and 1 year-desiccated samples.

### 3.5. Evaluation of *Coelastrella* sp. D14 for wastewater treatment

After changing carbon and nitrogen sources in BG11 medium, the growth of *Coelastrella* sp. D14 in wastewater was evaluated, more specifically in a piggery effluent. The composition of the piggery effluent used in the trials is shown in Table 1.

*Coelastrella* sp. D14 was submitted to an initial screening in different PWW concentrations (Fig. 9). Experiments demonstrated that D14 could grow in this effluent at concentrations up to 30% (v/v). However, the initial lag phase becomes longer with increasing PWW concentration. At 40% PWW, no visible growth was observed during the 15 days of the experiment. Native microorganisms could be found in the medium and co-cultivated with the inoculated D14. Nevertheless, in all experiments the inoculated microalga rapidly outcompeted other organisms present in the wastewater. Thus, their impact on growth and biomass concentration was considered insignificant.

**Fig. 9.**
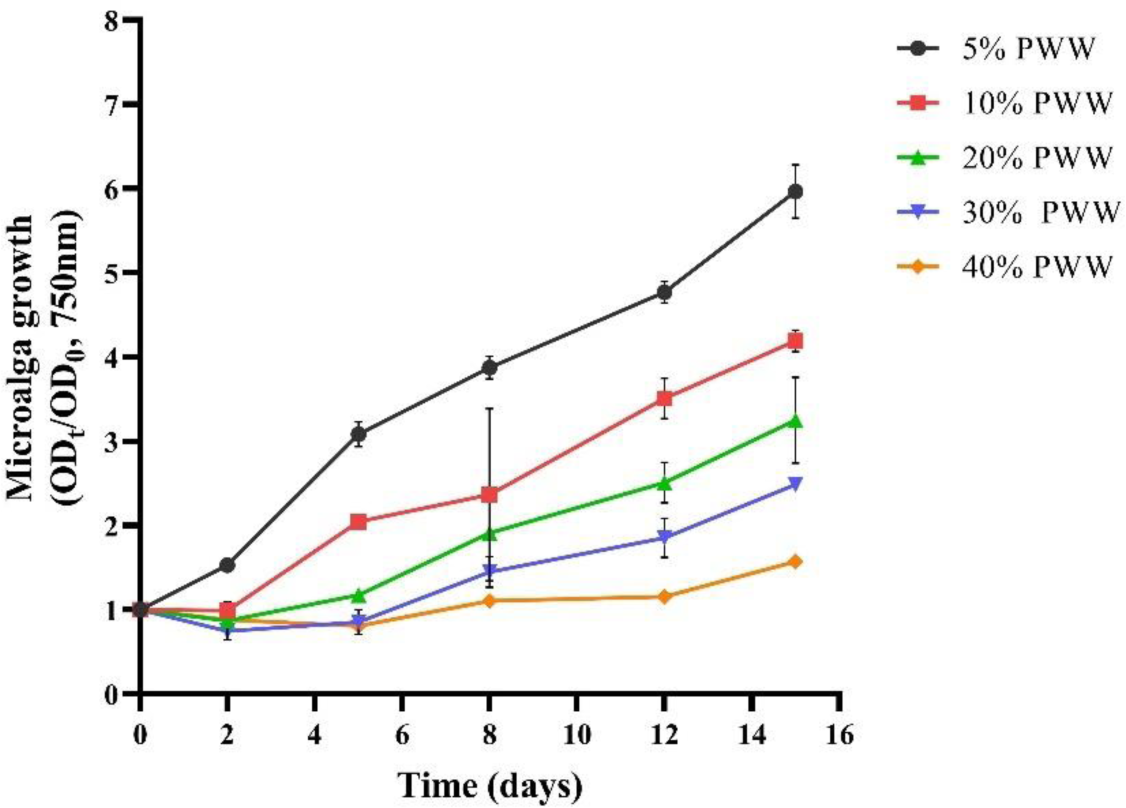
Growth of *Coestrella* sp. D14 at different concentrations of piggery wastewater (5, 10, 20, 30, and 40% v/v). In all cases, the graphs show the normalized value of the average OD_750nm_ together with the standard deviation (n=2).

*Coelastrella* sp. D14 grown in synthetic medium (BG11) presents a high amount of carbohydrates (37.8%), followed by 23.2% of protein, 13.9% lipids, and 7.1% ash. When grown in PWW, it has a similar composition profile for biomass grown in 5 and 10% PWW (30-34% carbohydrates, ∼21% protein, and 14-16% lipids).

### 3.6. Effect of *Coelastrella* sp. D14 as biostimulant for seed germination

Plant growth is affected by phytohormones, amino acids and polysaccharides, along with other nutrients, available from various sources, including microalgae. Here, the effect of *Coelastrella* sp. D14 biomass was evaluated on germination of cress seeds (*Lepidium sativum*). In the biostimulant assay, a germination index (GI) of 100% was attributed to distilled water (control). Values higher than the control were considered to have a biostimulant activity (Fig. 10). The highest GI values obtained were 128% corresponding to non-disrupted biomass trials at 1 or 2 g/L on BG11 medium and 132% at 1g/L culture on 5% PWW. Cell disruption causes a significant drop, for instance, at 2 g/L of D14 grown on BG11, yield dropped a 45% of the GI value with respect to the whole biomass.

**Fig. 10.**
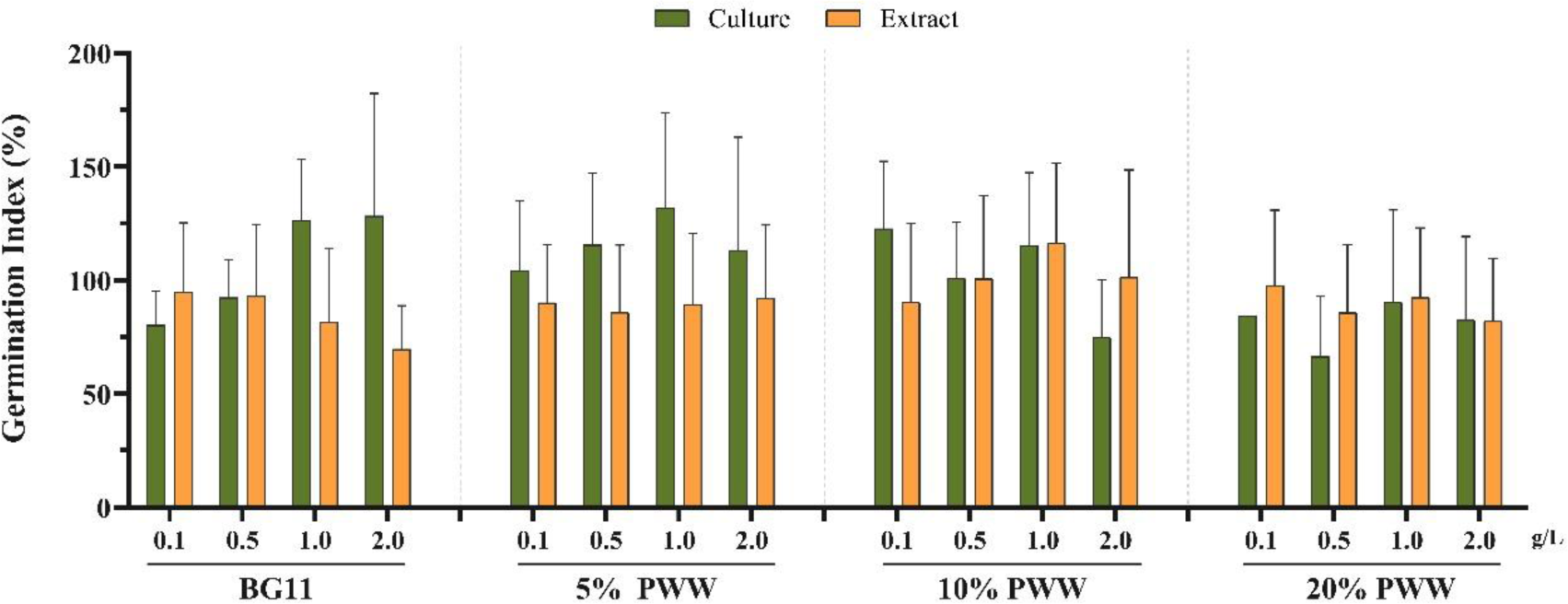
Germination index (%) of *Lepidium sativum* seeds using of *Coelastrella* sp. D14. The graph shows the mean of germination index (GI) value, either using initial biomass or an extract of D14 growth in four different media (BG11 or BG11 containing different concentrations of piggery wastewater, PWW). GI value obtained when using distilled water as medium was considered as control (GI, 100%). Error bars indicate, standard deviation (n = 20).

This tendency is not seen when grown on 20% PWW. In this case, there are not so many differences between both culture and extracts GI values, not reaching in any case the GI 100% value. In general, values under 100% may suggest that the microalga concentration or their biochemical composition might be excessive or toxic for *L. sativum*, negatively affecting their growth (Navarro-López et al., 2020).

Therefore, these results highlighted that 1 g/ L of whole algal suspension grown at 5% PWW may be the best treatment for root lengths. However, whole D14 biomass grown at 10% PWW and used at 0.1%, also yielded a GI value of approximately 123%. The latter is more interesting since it uses a higher percentage of PWW to grow, lower amount of non-processed biomass, decreasing also downstream costs.

## 4. DISCUSSION

The isolate obtained from the solar panel was identified as a *Coelastrella* sp. based on morphological features and phylogenetic analysis. The morphological features of the studied strain (single oval cells with a mean diameter of 8.68 ± 1.96 μm) match those of the genus *Coelastrella* which is characterized as unicellular and occasionally forming aggregates with vegetative cells spherical to subspherical, from 4.2–14.8 µm in diameter (Shetty et al., 2021; Doppler et al., 2022). The phylogenetic tree was obtained using conventional loci such the nuclear 18S rRNA. This novel strain belongs to the Chlorophyceae class, Sphaeropleales order, Scenedesmaceae family, *Coelastrella* genus. The closest relatives of D14 are *Coelastrella* sp. isolate 3A (*Coelastrella thermophila* var. globulina) (Boutarfa et al., 2022) and the unclassified *Coelastrella* sp. SAG 2471(GenBank: KM020087). The clade to whom D14 belongs contains in general *Coelastrella* strains isolated from harsh environments (Ki4-from a Japanese asphalt surface in midsummer, (Kawasaki et al., 2020); *Scenedesmus* sp. JB11 from an extreme saline-alkali soil, data from NICB; *Coelastrella* sp. isolate 3A isolated from an algerian hot spring, (Boutarfa et al., 2022)) and some of them with remarkable properties such as for producing n-6 and n-3 PUFA fatty acids (Boutarfa et al., 2022) or astaxanthin production (Kawasaki et al., 2020).

In the conditions tested, D14 displayed a doubling time of 2.6 days (Table S1). A novel freshwater *Coelastrella* strain isolated in Belgium, presented a rapid growth in phototrophy, with a doubling-time of 6.8 ± 0.30 h hours at a light intensity of 400 µmol·m^−2^·s^−1^ and 5% CO_2_ (Corato et al., 2022). The specific growth rate seems to depend on dosing times of the carbonic solution added to the culture unless indoor. This causes an increase on the lipids and proteins content with major carbon source dosing times, while the carbohydrate content decreased, suggesting that the carbon source is a critical parameter for algal growth (Razooki et al., 2019). The doubling time of D14 is given without any extra C source, which could explain the low value obtained and leaves room for improvement by increasing the % of CO_2_ or adjusting the amount of light during growth.

D14 growth was assayed under several conditions as microalgal growth and biomass production depends on nutrient availability and amounts, light intensities, pH or temperature value among others. Most microalgae can grow over the range of pH values from 6.8 to 8.0, the suitable pH value depending on the microalgal species (Daneshvar et al., 2021). Concretely, a range from 5.0 to 9.0 is reported for *Coelastrella* KKU-P1 strain (Thepsuthammarat et al., 2023). D14 grown in flasks is viable from a wide range of pH, from 4.0 to 11.0, although the highest production was obtained under a pH of 7.5 (Fig. 5). *Coelastrella* sp. strain D3–1 has been reported to resist from pH 2 to pH 11, but the experiments were performed in different conditions to this work, mainly, after the stress treatment on a diluted medium 0.2XBG11, cells were spotted on BG11 plates and grown for 7 days (Saito et al., 2023) while in this work data was taken in liquid medium with the pH corrected. Regarding the temperature, it greatly affected the growth of *Coelastrella* sp. D14 (Fig. S2) as no growth was observed over 50 °C. On the contrary, several *Coelastrella* have been reported to be able to grow on 50°C or over, for instance, *Coelastrella* sp. M60 (Nayana et al., 2022) or *Coelastrella* sp. strain D3–1 (Saito et al., 2023) resisted temperatures of up to 50 °C. However, D14 was able to resist desiccation up to one year (the maximum time tried). There are no similar experiments done with other *Coelastrella*. However, a heat-dry stress done in *Coelastrella* sp. D3–1, in which cell pellet was exposed to 42 ◦C for 3 h in a dryer, showed that the microalga was able to grow when later was spotted on BG11 plates (Saito et al., 2023). The xerotolerance that D14 showed is consistent with its original habitat, a solar panel, in which the water activity is logically low. It is quite possible that other *Coelastrella* isolated from solid environments such that characterized from an asphalt surface (Kawasaki et al., 2020) could display a similar behavior.

To enhance the biomass production, the growth of *Coelastrella* sp. D14 was investigated under both heterotrophic and mixotrophic conditions. The microalga was able to grow well heterotrophically using glucose, fructose, and mannose as carbon sources, hardly with sucrose and could not grow on maltose or lactose (Fig. 4A). In mixotrophic conditions, the growth of D14 reached the highest OD_750nm_ when the medium was supplemented with glucose, in comparison to the use of mannose or phototrophic growth. In general, green microalgae can efficiently use glucose and fructose for growing but they usually lack sucrose transporter systems (Pang et al., 2019). However, there are some reports as *Coelastrella* sp. KKU-P1 in which it is capable of sucrose consumption and could be used for growing on unhydrolyzed molasses as a low-cost carbon source that is rich in sugars, mainly sucrose, glucose, and fructose (Thepsuthammarat et al., 2023). D14 could be subjected to adaptive laboratory evolution experiments in a future to improve its mild growth on sucrose and in this way, being able to grow on more low-cost substrates.

In autotrophic conditions, the absence of nitrogen in the medium, D14 was not able to survive (Fig. 7). This is expected as nitrogen is an essential component for microalgae growth, needed for macromolecules synthesis among other things and *Coelastrella* is not able to fix it, at least in the conditions tested. When NaNO_3_ from BG11 was replaced by urea or NH_4_Cl (16 mM), growth was restored, although it was not as good as using the original NaNO_3_of BG11. Ammonium is the preferred nitrogen source for algae since it consumes less energy, as it does not require a redox reaction (Nayana et al., 2022).

Furthermore, the use of other cheap sources of nitrogen as urea, is convenient for reducing economical costs of the microalgae growth. Due to the ability of microalgae to grow in very diverse environments, and considering the idea of circular economy, wastewaters that are rich in nutrients can be used as a culture medium. In fact, growing microalgae in wastewater is a suitable alternative to reduce freshwater expenses and valorize residual nutrients (Ahmed et al., 2022; Sánchez-Quintero et al., 2023). (Sharma et al., 2022) enumerates several economic and growth considerations when cultivating microalgae in wastewater pointing to that appropriate strains’ selection is crucial for the whole process. Therefore, it is important to count on different strains or consortiums able to bioremediate the effluents for choosing the best one for a certain valorization, for instance, using the produced biomass as plant biostimulant.

One kind of wastewater that causes big concern is the one from piggery industry (Ferreira et al., 2021), which is a complex effluent rich in nutrients, such as ammonia and organic matter. It causes eutrophication and toxicity of freshwater ecosystems while the deep dark color hampering photosynthesis of this medium when discharged into rivers without complete treatment (Li et al., 2019; Ferreira et al., 2021; Lee et al., 2021). On the other hand, when using piggery effluents directly in composting for agriculture, greenhouse gas emissions are generated (such as CO_2_ and N_2_O) (Mohedano et al., 2019; Hu et al., 2020). For this reason, several attempts have been reported for the biological treatment of raw PWW with microalgae with different approaches. The studies at a pilot scale treating undiluted raw PWW with microalgae, also resulted in an efficient removal of nutrients and an enhancement in the clarity of wastewater (Lee et al., 2022). Another *Coelastrella* sp. isolated from an ammonia-rich environment was used for PWW treatment in a 4-day two-step process: heterotrophic plus mixotrophic steps in a narrow transparent photobioreactor (Lee et al., 2021). In these conditions, *Coelastrella* sp. could remove 99% of ammonia, 92% of chemical oxygen demand (COD), and 100% of phosphorus. In this case, the microalgal biomass was oriented towards the production of biodiesel of high quality (Su et al., 2023). This study showed that D14 can grow on piggery wastewater up to 20%. More studies must be done to evaluate how the microalga biochemical composition could be affected.

In this work, besides assessing the potential of *Coelastrella* sp. D14 to grow in piggery wastewater, it was explored, the use of the resulting biomass to stimulate plant growth. A bioassay was performed based on the germination index of *Lepidium sativum* seeds. The results have shown that D14 has potential as a biostimulant product acting as a gibberellin-like when growing on BG11 at 1-2 g/L or 5% PWW at 1 g/L, yielding GI values up to 132%. Other bioassays in agricultural models, such as examining root formation in mung beans and cucumbers, will be necessary to further demonstrate the auxin-like effect. On the other hand, data obtained with the whole biomass yielded higher GI values with respect to the broken cells. For other strains such as the microalga *Scenedesmus obliquus* a similar behavior has been reported: the highest GIs were obtained with the initial biomass in the absence of any pre-treatment (Navarro-López et al., 2020). For this microalga, grown on brewery effluents, a GI of 139% was obtained.

There are recent strategies of culturing microalgae using wastewater and CO_2_ to produce large quantities of biomass at moderate costs while integrating local and circular economy approaches (Sánchez-Quintero et al., 2023). The fact that the microalgal D14 biomass could be used directly as a biostimulant implies a reduction in economic costs and a sustainable application, avoiding synthetic stimulants. Regarding the use of raw piggery wastewater, despite that microalgal treatment of reduces the risk of pathogens by 63% (Lee et al., 2022), recent reports highlight their persistence. To align with EU Regulation 2019/1009 on plant biostimulants, it is recommended to use extracts to mitigate potential pathogen presence (Lee et al., 2022; Sánchez-Quintero et al., 2023). This needs for optimization studies on the preparation of alga extracts, as stated procedures may impact the final bioavailability of microalga compounds. These factors are influenced by the microalga species, cultivation medium, and culture state (Ferreira et al., 2018; Ferreira et al., 2019). More specifically, bioactive compounds reported as biostimulants, such as phytohormones, heteropolysaccharides, amino acids, or vitamins (as reviewed by (Sánchez-Quintero et al., 2023)), are produced in various phases of growth (Tan et al., 2021). This variability could explain why extracts with the same concentration but grown in different media (resulting in distinct growth curves) do not exhibit the same biostimulant behaviour. For this reason, it could be convenient in the next future to have more studies on *Coelastrella* sp. D14 broken cells at different conditions to optimize its use as biostimulant.

## 5. Conclusion

Microalgae are a promising feedstock to produce valuable products, but it is important to isolate strains with the ability to grow in stressful conditions, to avoid contaminations and having a particular biochemical profile, and to widen their biotechnological applicability. This research highlights the potential of the strain *Coelastrella* sp. D14 for growing on low-cost resources such as PWW while removing nutrients from this effluent. If grown on 5% PWW, D14 biomass could also be used as a biostimulant allowing a more sustainable process.

## Author contributions

All authors have contributed to the manuscript. SB and GG contributed equally to the conception and design of the study. SB, GG, and AF conducted the characterization and studies on the strain isolates and performed the studies. All authors analyzed and discussed the data. SB, GG and JMN wrote the draft of the manuscript, and AF and LG made the revisions. All authors read and approved the submitted version.

## CRediT author contribution statement

**Sara Baldanta**: Conceptualization, Investigation, Methodology, Software, Writing-Original draft preparation. **Alice Ferreira**: Investigation, Software, Visualization, Writing-Original draft preparation. **Luisa Gouveia**: Conceptualization, Visualization, Supervision, Resources, Writing-Reviewing and Editing. **Juana Maria Navarro Llorens**: Writing-Reviewing and Editing, Resources, Supervision. **Govinda Guevara**: Conceptualization, Investigation, Data curation, Writing-Reviewing and Editing.

## Data availability

The 18S rRNA data has been submitted on Genbank under accession number PP158241. Data will be made available on request.

## Acknowledgments

Authors would like to thank José Luis García and Beatriz Galán (CIB-CSIC) for their technical support and Raquel Arnal (UCM) for her help in some of the studies. We are also grateful to the “Programa Iberoamericano de Ciencia y Tecnología para el Desarrollo” (CYTED) through RED RENUWAL 320rt0005. The authors would like to thank Valorgado for supplying the piggery wastewater, Graça Gomes, and Natércia Sousa (LNEG) for laboratorial assistance and maintenance of the microalgae cultures.

## Supplementary material

**Table S1.**
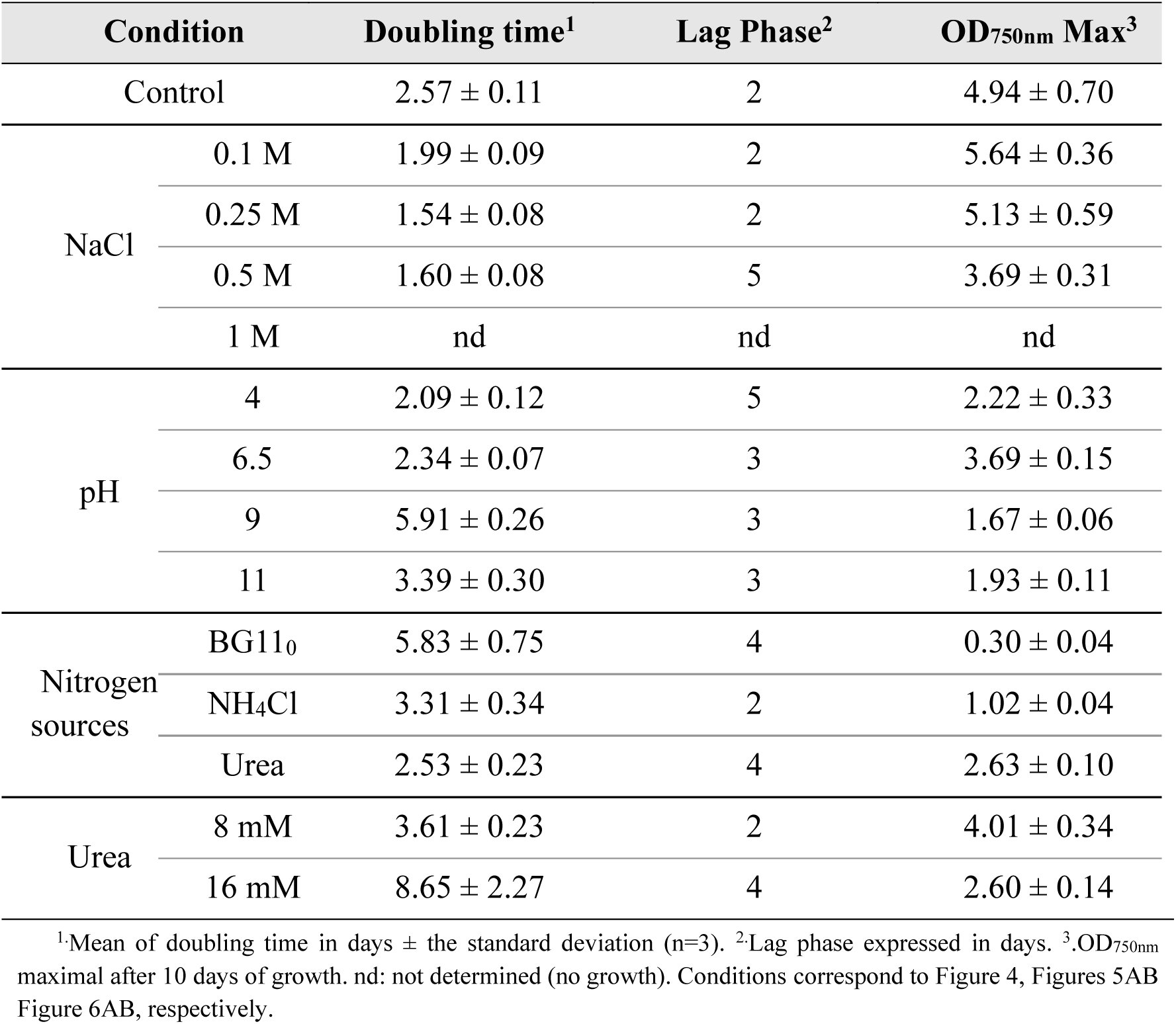
Growth parameters of *Coelastrella* sp. D14 growing in different conditions.

**Fig. S1.**
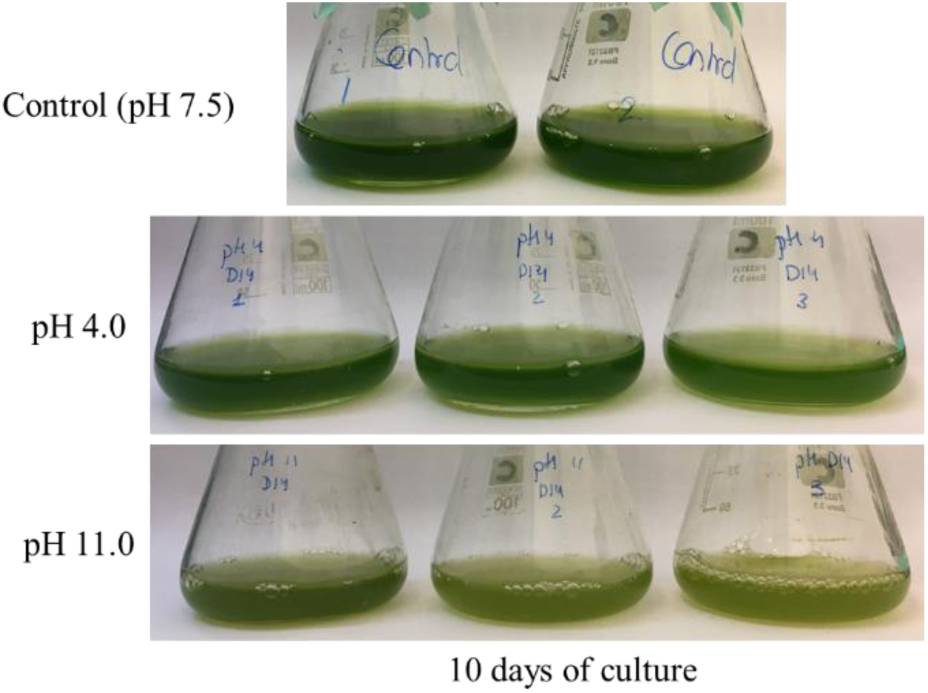
Growth of *Coelastrella* sp. D14 at different pH.

**Fig. S2.**
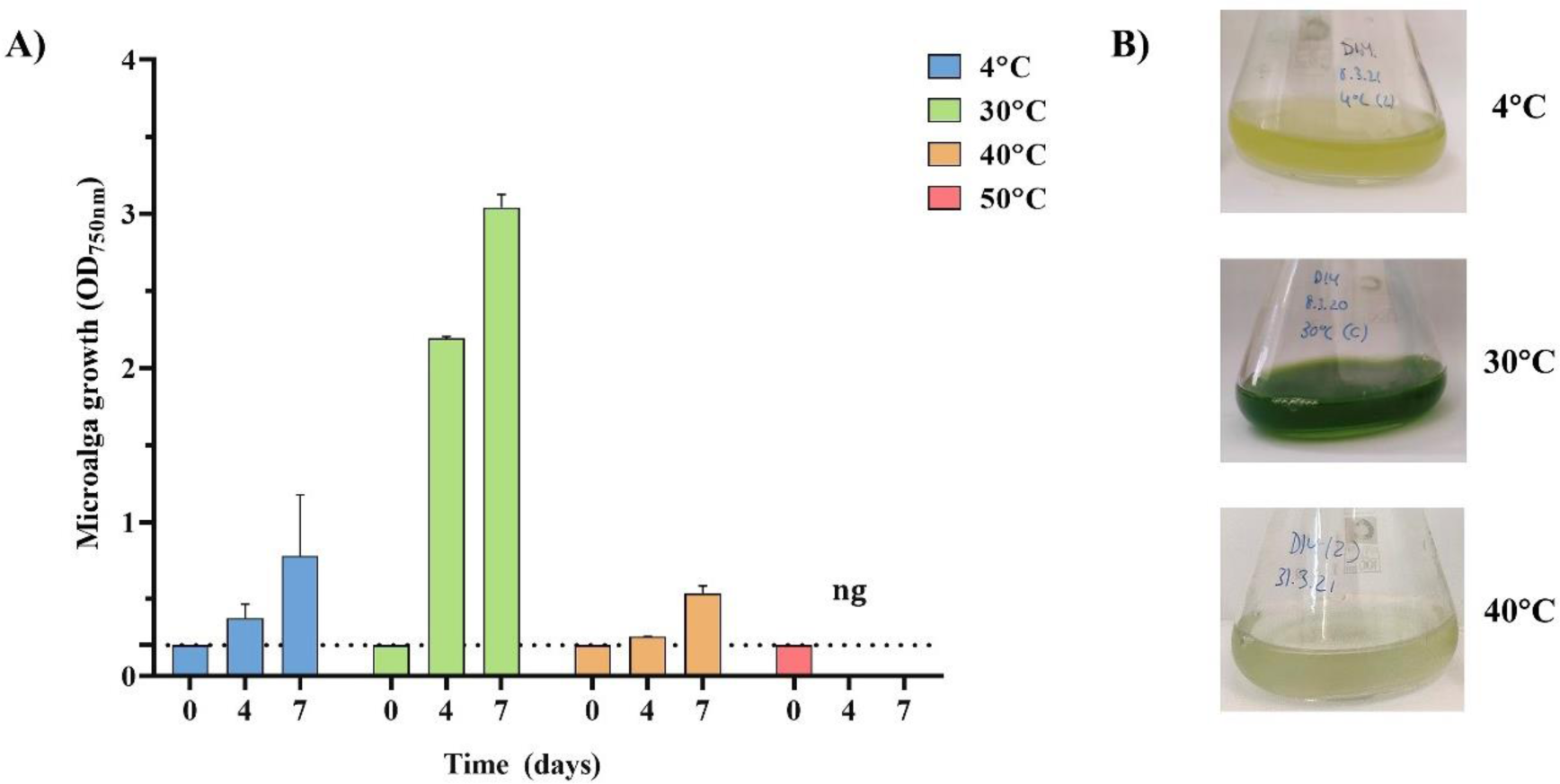
Growth of *Coelastrella* sp. D14 at different temperatures. A. OD_750nm_ reached after 10 days of growth. Initial OD_750nm_ was 0.2 (dashed line). B. Pictures taken from different points of the experiment. ng: no growth.

